# Comparative metagenomics of coalbed methane microbial communities reveals biogenic methane potential in the Appalachian Basin

**DOI:** 10.1101/319590

**Authors:** Daniel E. Ross, Daniel Lipus, Kelvin B. Gregory, Djuna Gulliver

## Abstract

Natural gas is a major source of global energy, and a large fraction is generated in subsurface coalbed deposits. Microbial communities within coalbed deposits impact methane production, and as a result contribute to global carbon cycling. The process of biogenic coal-to-methane conversion is not well understood. Here we demonstrate the first read- and assembly-based metagenome profiling of coal-associated formation waters, resulting in the recovery of over 40 metagenome-assembled genomes (MAGs) from eight individual coalbed methane wells in the Appalachian Basin. The majority of samples contained hydrogenotrophic methanogens, which were present in higher relative abundances than was previously reported for other coalbed basins. The abundance of Archaea and salinity were positively correlated, suggesting that salinity may be a controlling factor for biogenic coalbed methane. Low-abundance coalbed microbial populations were functionally diverse, while the most dominant organisms exhibit a high degree of genomic and functional similarities. Basin-specific pan-metagenome clustering suggests lower abundant and diverse bacterial communities are shaped by local basin parameters. Our analyses show Appalachian Basin coalbed microbial communities encode for the potential to convert coal into methane, which may be used as an indicator of potential biogenic methane production for future well performance and increased well longevity.

---

Methane is an important fossil energy resource in the global energy landscape. As conventional natural gas resources become depleted, unconventional gas technologies will emerge with greater importance for global energy security. One unconventional gas technology is coalbed methane (CBM), which relies on underutilized natural methane repositories trapped in subsurface coalbeds. CBM wells provide access to subsurface methane deposits at a reduced cost and environmental impact relative to traditional mining practices ^1^.

An estimated 40% of U.S.-based CBM is biogenic^2,3^, and recently, there has been a growing interest in understanding microbial communities in coalbed deposits, as these communities may be potential indicators of productive CBM wells or utilized to enhance *in situ* production of methane.^4^ Research in this area has offered insight into unique subsurface microbial pathways that affect methane production and ultimately, the global energy supply and global carbon cycling. Biological conversion of coal to methane is complex and requires multiple enzymatic steps, performed by a diverse set of microorganisms, including hydrocarbon degraders and methanogens^5^—unraveling these pathways, and the associated diverse community of microorganisms, is paramount to understanding biogenic methane production.

Much of what is known about coalbed microbial communities has been derived from 16S rRNA gene sequencing^6–8^, which provides important information about microbial community structure and diversity, but lacks functional data required for detailed metagenomic analyses. A limited number of studies have provided valuable insight into the metabolic potential of CBM systems using metagenomics^9,10^. However, these studies are limited to the western (Alberta Basin, Powder River Basin, and the San Juan Basin) and interior basins (Illinois Basin), neglecting large coalbed basins in the eastern United States, specifically the Appalachian Basin. The Appalachian Basin is an energy rich expanse that stretches from New York to Alabama and according to the newest EIA report is one of the most productive energy regions in the nation (www.eia.gov).

To characterize the unknown microbial community in the eastern US coal basin and gain insight in its potential for biogenic methane production, we present the first metagenomic investigation of Appalachian Basin coalbeds and the first coalbed-focused pan-genomic comparison across geographically distinct regions. Findings from this study provide 1) a read-based and assembly-centric taxonomic profiling of the previously uncharacterized microbial communities, 2) the first Appalachian Basin coalbed metagenome-assembled genomes (MAGs), 3) functional potential characterization of coalbed microbial communities extant in the Appalachian Basin, and 4) a taxonomic and functional comparison of Appalachian Basin metagenomes to microbial communities from geographically distinct coalbeds. Our results demonstrate the Appalachian Basin to be distinct from previously characterized basins, specifically an abundance of hydrogenotrophic methanogens from the Order *Methanomicrobiales*, suggesting a novel subsurface environment that could be targeted for methane production and contributes to unknown amounts of carbon cycling. We show abundant populations were highly conserved across geographically distinct coalbed basins while low-abundance microorganisms were functionally diverse within and across coal basins. This work provides a framework for understanding subsurface microbial metabolisms and the potential for coal to methane conversion, which can be utilized as an indicator for the potential to recover methane from unmined areas, and enable the development for microbial enhanced coalbed methane strategies in the Appalachian Basin.

## METHODS

### Sample collection and chemical characterization

Formation water samples (*i.e*., water from CBM wells produced after drilling) were collected from eight separate coalbed methane wells from the Appalachian Basin. Samples were extracted from CBM wells named Key8, Key7, K34, K35, P21, BB137, MC79, and L32A, at a depth of 796 ft., 1,201 ft., 1,704 ft., 1,912 ft., 1,961 ft., 1,980 ft., 2,239 ft., and 2,578 ft., respectively. All samples were obtained from the Pocahontas 3 coal seam in December 2015 (Buchanan County, VA; Tazewell County, VA; McDowell County, WV) with the exception of Key8 (Middle Kittanning) and Key7 (Upper Kittanning/Brookville "A”), which came from Indiana County, PA and Westmoreland County, PA, respectively. Formation water samples were collected in sterile 1L bottles, placed on ice (immediately following extraction from the well), and frozen upon arrival at NETL. The coal rank associated with production water ranged from low/medium volatile bituminous (K34, K35, P21, BB137, MC79, and L32A) to medium/high volatile bituminous (Key7 and Key8). Cumulative Appalachian Basin coalbed methane production ranged from 10 billion cubic feet (BCF) to 1 trillion cubic feet^11^.

Major and trace cations and anions (Na, Ca, Mg, K, Fe, Sr, Ba, Li, Mn, and Cl) were examined at the Pittsburgh Analytical Laboratory (National Energy Technology Laboratory) following EPA methods 6010C and 300.1 using ICP-OES (Perkin Elmer Optima 7300 DV) and IC (Dionex Ion Chromatograph). Samples were run in duplicate, with average values being recorded (Supplementary Table 1).

### Read-based microbial abundance correlation

Read-based taxonomy (Archaea and Bacteria relative abundance) was correlated to major and minor cation/anion concentrations with Spearman rank coefficients using vegan in R^12,13^.

### DNA extraction, metagenome sequencing, and assembly

Aliquots of formation water (50 mL) were filtered through a 0.2 μm filter and DNA was extracted from the filter using the MoBio PowerSoil or PowerWater DNA Isolation Kits (Carlsbad, CA). For shotgun metagenome sequencing, DNA libraries were prepared using the Nextera XT DNA Library Preparation Kit according to manufacturer's protocol (Illumina, San Diego, CA). Paired-end sequencing reads (2 × 300 bp) were generated on an Illumina MiSeq with the MiSeq v3 Reagent Kit (600 cycles) (Illumina, San Diego, CA). Initial assessment of reads was performed with Kaiju using default settings: *nr* reference database, no SEG low complexity filter, greedy run mode, minimum match length = 11, minimum match score = 70, and allowed mismatches = 5^14^; MG-RAST^15^; GraftM (https://github.com/geronimp/graftM); and Phylopythia (Supplementary Table 2)^16^. Paired end reads were assembled using metaSPAdes version 3.8.0^17^ with error correction and with k-mer values of 33, 55, and 77. Assembly quality metrics (*e.g*., GC-content, N50, number of contigs, longest contig) were assessed using QUAST^18^ (Supplementary Table 3; Supplementary Figure 1). Percentage of reads mapping to metagenome contigs was performed with the ‘map reads to contigs’ tool in the CLC Genomics Workbench version 8.5.1 (CLC Bio, Aarhus, Denmark). Annotation of assembled metagenome contigs was performed with PROKKA^19^ and MG-RAST^15^.

Contigs generated from metagenome assembly were binned using reference-independent methods (VizBin^20^). Completeness of metagenome-assembled genomes (MAGs) was assessed with CheckM, using single-copy marker genes^21^. The lineage_wf was initially employed, and combined with BLASTn and Phylopythia outputs to further interrogate MAG quality using the user-defined taxonomy_wf. For example, bins defined at the phylum level (*e.g., Euryarchaeota*) using the lineage_wf were analyzed further and based upon sequence similarity to *Methanocalculus* or *Methanoregula*, order-level taxonomy_wf was used to assess completeness and contamination using the *Methanomicrobiales* marker gene set. MAGs generated from contig binning were annotated using the RAST annotation server^22^ and PROKKA^19^. Manual curation of MAGs was performed using output from CheckM and RAST. Quality designation for metagenome assembled genomes (MAGs) or population genomes was based upon metagenome-assembled genome (MIMAG)^23^ standards developed by the Genomic Standards Consortium (GSC), with high-quality draft genomes having >90% completion, <5% contamination, with 23S, 16S, and 5S rRNA genes and >18 tRNAs. Medium-quality draft genomes were characterized by having >50% completion, and <10% contamination, while low-quality draft genomes were <50% complete and <10% contamination.

DNA sequences for metagenome samples from the San Juan basin (CG7, CG8, CG13, and CG19) and Alberta Basin (CO182, CO183, and Trident1560D) were downloaded from the Hydrocarbon Metagenomics Project (HMP) database (http://hmp.ucalgary.ca/HMP/). Full details of sample collection, DNA extraction, and sequencing can be found online (http://hmp.ucalgary.ca/HMP/) and has been documented by An and coworkers^9^. Raw read statistics are shown in Supplementary Table 4 and 5. Metagenomes can also be found at the metagenomics analysis server (MG-RAST) under the Hydrocarbon Metagenomics Project (mgp2360). Produced water samples CG7, CG8, CG13, and CG19 from the San Juan Basin, and Trident1560D from the Alberta Basin were sequenced using 454-pyrosequencing. Samples CO182 and CO183 from the Alberta Basin were sequenced with Illumina.

Average nucleotide identity (ANI) and average amino acid identity (AAI) were calculated with the enveomics collection toolbox^24,25^.

### Pan-metagenome analysis

Pan-metagenomics was performed with the Bacterial Pan Genome Analysis Tool (BPGA) version 1.0.0^26^. Default BPGA parameters were employed, which included USEARCH clustering at 50% sequence identity cutoff^27^.

### Data availability

The datasets generated during and/or analyzed during the current study are available on the Energy Data Exchange (EDX) page (https://edx.netl.doe.gov/dataset/appalachian-basin-metagenome-sequencing-reads).

## RESULTS

### Read-based taxonomy and functional profiling of Appalachian Basin formation water metagenomes

Formation water collected from eight separate coalbed methane wells was utilized for metagenomic analysis and geochemical characterization. Taxonomic profiling of metagenome reads revealed that Appalachian Basin metagenomes contained varied proportions of Archaea and Bacteria, with the relative abundance ranging from 1–81%, and 19–99%, respectively (Figure 1). The relative abundance of Archaea was >48% for five of the eight metagenomes, greater than what has been found in other coalbed basins^6–8^, but similar to Archaea abundances in crude oil reservoirs^28^.

**Figure 1.**
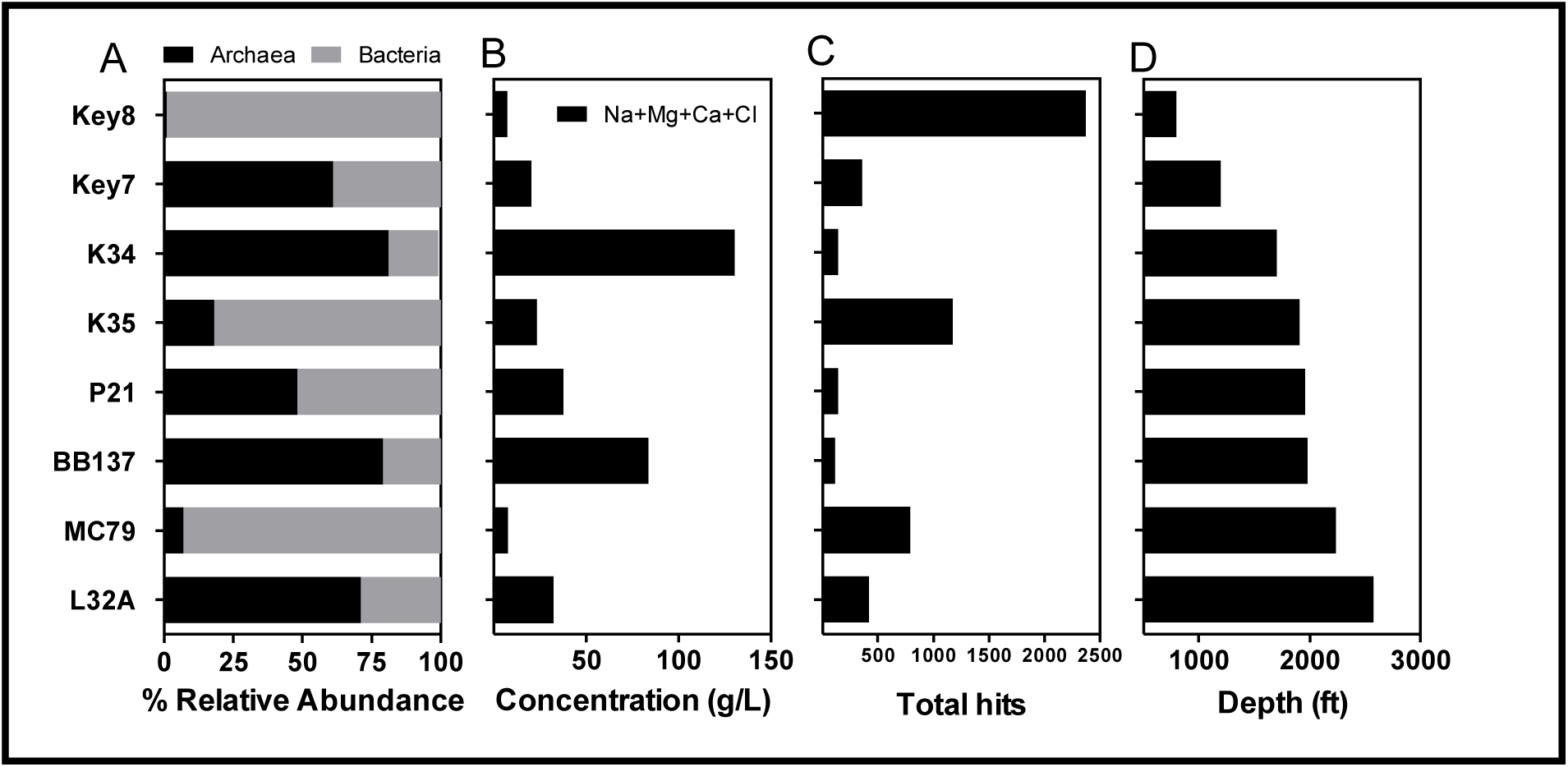
Read-based taxonomy profiling, and geochemistry of eight Appalachian Basin formation water samples. A) Class-level taxonomy of coalbed methane metagenomes from Illumina sequencing reads. B) Major ions found in formation water samples. C) Total number of reads mapping to hydrocarbon degradation pathways. D) Depth profile of formation water samples.

Environmental factors, such as salinity, coal rank, and depth, are known to play a key role in shaping the taxonomic distribution of subsurface microbial communities^10,29–31^. Coalbed-associated formation water geochemistry was examined and varied from moderately to highly saline, with concentrations of major ions (Na, Cl, Mg, and Ca) ranging from 7.29 g/L (Key8) to 130.30 g/L (K34), (Figure 1; Supplementary Table 2). There was a positive correlation between Archaea abundances and salinity and a negative correlation between Bacteria abundances and salinity (Figure 1, Supplemental Table 2) in the Appalachian Basin samples. No observable trends were found between microbial community composition and associated coal rank (*i.e*., maturity) or depth.

Utilization of coal as a carbon and energy source requires breakdown of complex polymers into fermentable substrates, a poorly understood process and hypothesized to be the rate-limiting step in the conversion of coal to methane^5^. It is thought that hydrocarbon-activating organisms (*e.g*., Deltaproteobacteria and Firmicutes) are responsible for the initial attack on coal^32^. Examination of metagenome reads revealed the presence of hydrocarbon degradation pathways in all samples, though samples containing a higher abundance of Bacteria-specific sequences had a higher proportion of hydrocarbon degradation pathways (Figure 1; Supplementary File 1).

Samples containing an abundance of Bacteria-specific reads were predominantly comprised of Proteobacteria and despite phylum-level similarities across Appalachian Basin metagenomes, noteworthy distinctions were apparent at the class level. For example, metagenome L32A contained mostly Alphaproteobacteria (14%), metagenomes K35 and MC79 were comprised predominantly of Gammaproteobacteria (52% and 46% *Pseudomonas*, respectively), and Key8 was comprised predominantly of Deltaproteobacteria (22% *Desulfobacterium*) (Figure 2). Within the Proteobacteria phylum, Alphaproteobacteria and Deltaproteobacteria were most diverse across Appalachian Basin samples, while Betaproteobacteria, Gammaproteobacteria, and Epsilonproteobacteria were predominantly comprised of Burkholderiales, Pseudomonadales or Alteromonadales, and Campylobacterales, respectively. Four of the eight metagenomes containing the lowest relative abundances of Bacteria (<40%) had a higher proportion of members from the Terrabacteria group (Firmicutes, Chloroflexi, and Actinobacteria), FCB group (Bacteroidetes), and Synergistales.

**Figure 2.**
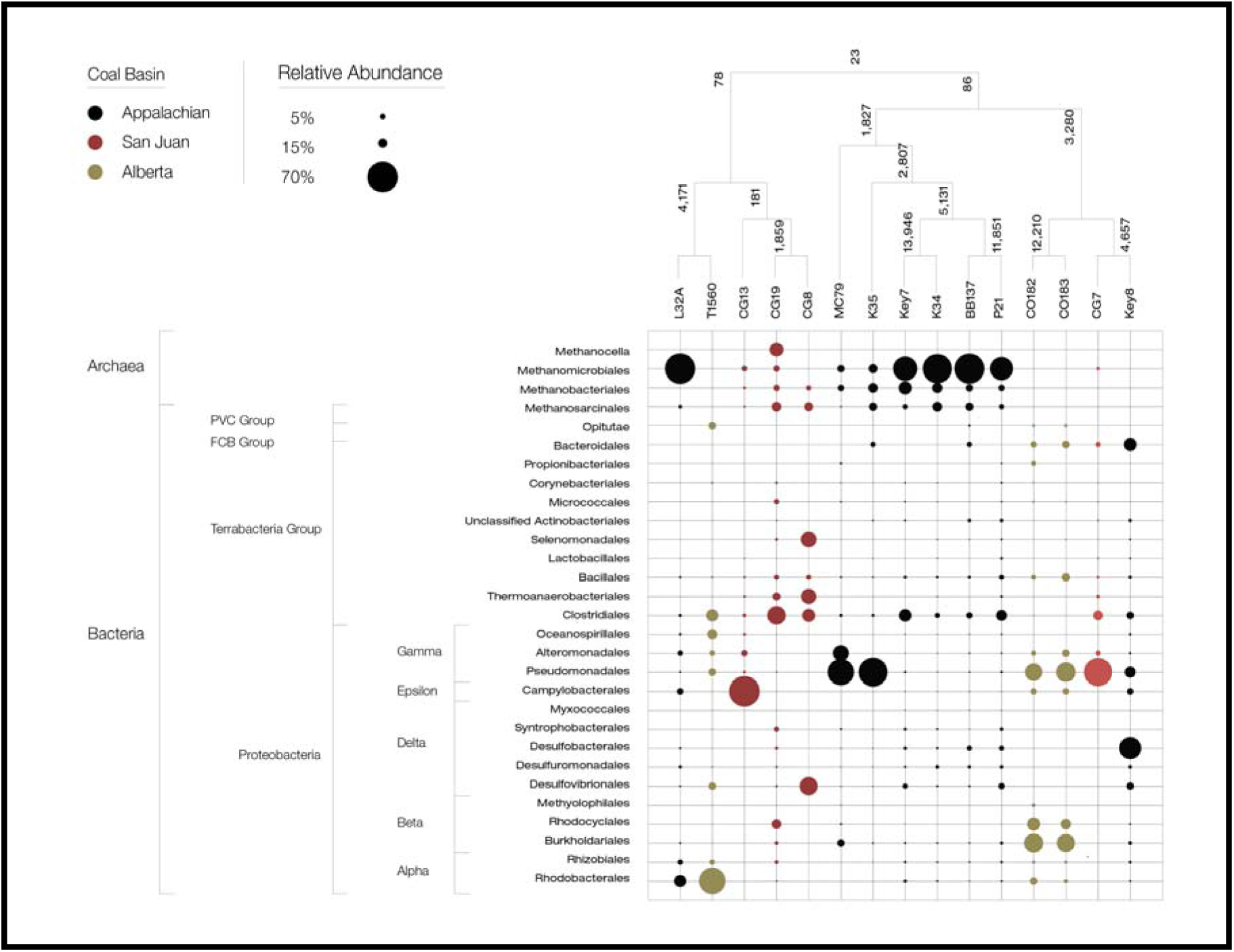
Assembly-based pan-metagenome analysis and read-based taxonomy of fifteen coalbed methane metagenomes from the Alberta Basin, Appalachian Basin, and San Juan Basin. Pan-metagenome clustering of coalbed methane metagenomes, with Bacteria and Archaea read distribution for Alberta Basin (yellow), Appalachian Basin (black), and San Juan Basin (red). Numbers on the pan-genome tree represent the number of gene families common across samples.

**Figure 3.**
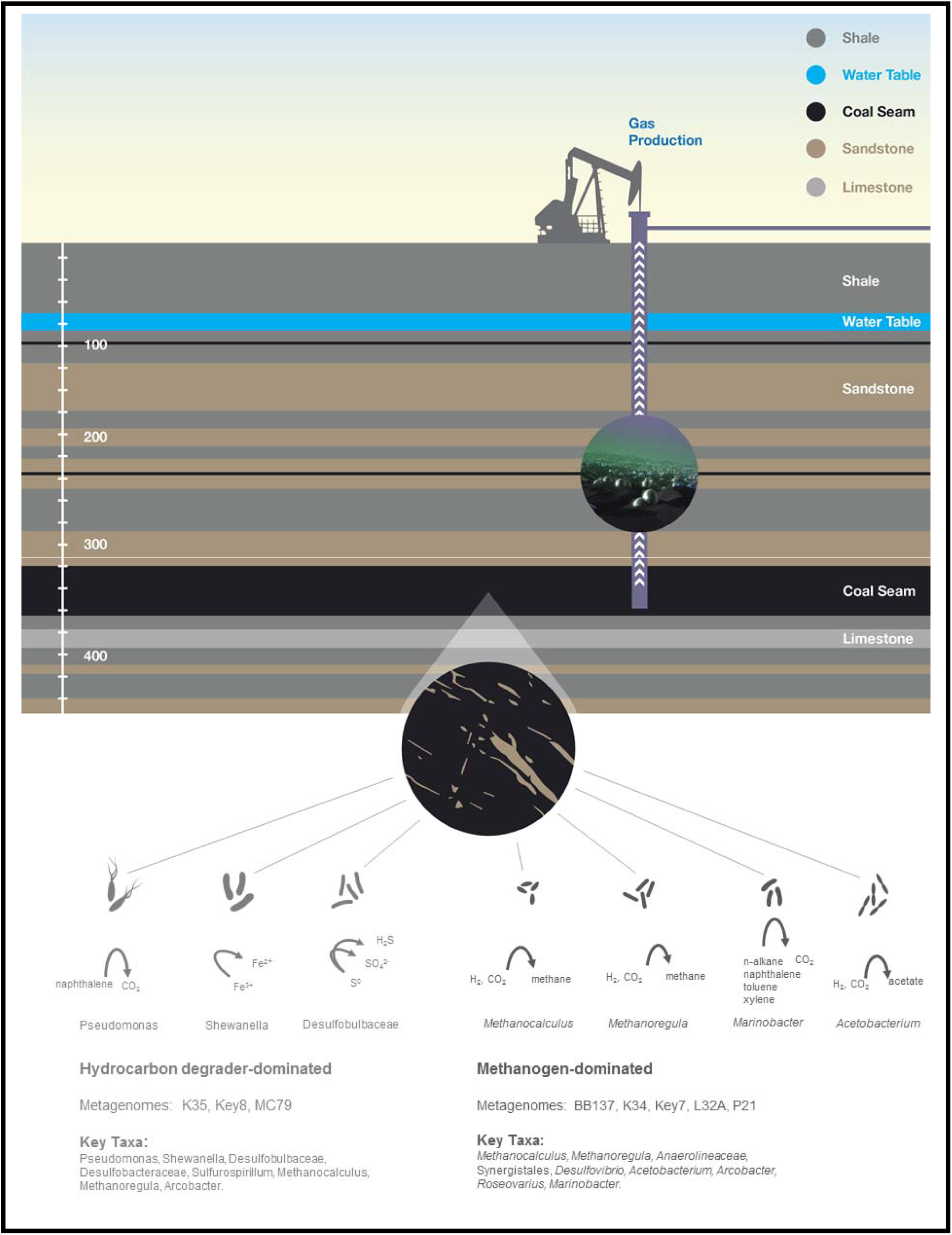
Metabolic overview and potential functional coal-to-methane conversion in coalbed methane systems. Metagenome data suggests two coalbed regimes; one dominated by hydrocarbon degraders, the other dominated by methanogens.

Closer examination of Archaea-specific reads revealed a predominance of Methanomicrobiales in the majority of Appalachian Basin samples, with relative abundances reaching 69% (Figure 2). Members of the order Methanomicrobiales are hydrogenotrophic methanogens that utilize H_2_/CO_2_ and formate, and inhabit a variety of environments, including marine and freshwater sediments^33^, and coalbed environments across the United States and abroad^6,7,34^. Other less-abundant Archaea orders represented in metagenome samples included Methanobacteriales and Methanosarcinales (Figure 2).

### Reconstruction of Appalachian Basin metagenome-assembled genomes (MAGs)

Assembly-based metagenome profiling was performed to obtain genome-resolved metabolisms of eight coalbed-associated microbial communities. Approximately 40 medium- to high-quality MAGs were recovered, with 77–94% of reads mapping to the metagenome assembled contigs. From the most abundant taxa we recovered ten Methanomicrobiales (*Methanocalculus* and *Methanoregula*), four Gammaproteobacteria (*Pseudomonas* and *Shewanella*), and two Deltaproteobacteria (*Desulfobacteraceae* and *Desulfobulbaceae*) population genomes (Supplementary Table 7).

#### Methanomicrobiales

Of the ten Methanomicrobiales MAGs, six were most closely related to halotolerant and hydrogenotrophic methanogens from hydrocarbon resource environments (oil reservoirs)—specifically, *Methanocalculus* sp. 52_23, *Methanomicrobiales* archaeon 53_18, and *Methanocalculus halotolerans* (Supplementary Figure 2-4). Four were most closely related to *Methanoregula formicicum* (Supplementary Table 8; Supplementary Figure 2-5).

Methanomicrobiales MAGs were highly similar, with ANI and AAI values >95%, suggesting that five of the six *Methanocalculus* MAGs belong to the same species, and three of the four *Methanoregula* MAGs belong to the same species (Supplementary Table 8). While samples contain acetoclastic methanogens, the predominance of Methanomicrobiales suggests hydrogenotrophic methanogenesis is operative in Appalachian Basin (Figure 2). Furthermore, acetoclastic methanogenesis may be inhibited by coal degradation products, which are similar in composition to crude oil, and contain substances that may interfere with this microbial metabolism^35^.

Following assembly and manual curation of *Methanomicrobiales* MAGs, comparative genomics was performed on the six *Methanocalculus* population genome bins (Supplementary Figure 5) to identify the functional similarities of the predominant methanogens across the Appalachian Basin. The core genome (predicted protein sequences common among all genomes examined) contained 1583 gene families (representing 44% of the total gene families) with 69% of the representative sequences most closely related to *Methanocalculus* sp. 52_23, 22% most closely related to *Methanomicrobiales*, 2.3% most closely related to *Methanoculleus*, and 1.4% most closely related to *Methanoregula*. Analysis of the *Methanocalculus* core genome revealed complete pathways for carbon fixation (reductive pentose phosphate cycle, and incomplete reductive citrate cycle), methane metabolism [methanogenesis (CO_2_ to methane, and acetate to methane), F420 biosynthesis, and acetyl-CoA pathway (CO_2_ to acetyl-CoA)], branched chain amino acid metabolism, ATP synthesis (V-type ATPase), and transport systems (molybdate, tungstate, glycine betaine/proline, osmoprotectant, phosphate, iron, zinc, cobalt/nickel, and lipooligosaccharide) (Supplementary File 2 and 3).

#### Gammaproteobacteria

Four MAGs related to Gammaproteobacteria were recovered from metagenomes K35 and Key8 (*Pseudomonas stutzeri*), and MC79 (*Shewanella putrefaciens*, and *Pseudomonas libanensis*)*. Pseudomonas* sp. K35 and *Pseudomonas* sp. Key8 are high-quality population genomes with a high degree of similarity to one another (ANI = 97.88%), and most closely related to *Pseudomonas stutzeri* CCUG29243^36^. Prior work has described a high abundance of *Pseudomonas* in coalbed systems^9,10,37,38^.

Formation water from wells Key8 and K35 were moderately saline (7–18 g/L). One mechanism of salinity tolerance employed by bacteria involves production of compatible solutes to maintain turgor pressure and water content within the cell^39^. *Pseudomonas* sp. K35 and *P*. sp. Key8 genomes contain the *ectABCD-ask* gene cluster, which encodes for biosynthesis of the compatible solute, hydroxyectoine^40^.

*Pseudomonas* may play an important role in coal solubilization as many *pseudomonads* have been reported to produce a variety of rhamnolipids— surface-active agents (surfactants) with the ability to solubilize hydrophobic carbon sources^41–43^. Production of rhamnolipids is controlled by many factors, including temperature, salinity, pH, and nutrient availability^43,44^ and may be induced by coal or coal substituents^42^. Both *Pseudomonas* sp. K35 and Key8 genomes contain the *rmlBDAC* operon for l-rhamnose biosynthesis (Supplemental Text), suggesting a potential for rhamnolipid-mediated coal solubilization in the Appalachian Basin. The solubilized coal constituents may be utilized directly by *Pseudomonas* or may provide fermentative substrates for other community members^5^.

Polycyclic aromatic hydrocarbons (PAHs) are a major constituent in coalbed production water and PAH-degrading microorganisms are prevalent in coal reservoirs^38,45^. *Pseudomonas* sp. K35 encodes for the upper and lower naphthalene degradation pathways and has the potential to degrade PAHs^36^. While we did not find naphthalene degradation pathways in *Pseudomonas* sp. Key8, other community members in the Key8 metagenome could perform this function, as the Key8 metagenome encodes for hydrocarbon degradation pathways (Supplementary File 1).

The other abundant Gammaproteobacteria genome bin was most closely related to *Shewanella*, with ~17% of metagenome reads mapping to the genome bin. The *Shewanella* population genome was 96% complete, and based on whole genome comparisons, was most closely related to *Shewanella putrefaciens* (ANI = 98.57%) (Supplementary Text, Supplementary Table 7). The genus *Shewanella* contains metabolically versatile facultative anaerobes best known for the dissimilatory reduction of metals (e.g. iron, manganese, and elemental sulfur)^46^ and *Shewanella* spp. have been isolated from coal environments^47^. The Appalachian Basin coalbed *Shewanella* population genome bin encoded for genes involved in assimilatory sulfate reduction, c-type cytochrome biogenesis (required for heme proteins involved in dissimilatory iron and manganese reduction), and nitrate reduction (Supplemental Text).

#### Deltaproteobacteria

Deltaproteobacteria were predominant in metagenome Key8, which contained two population genome bins related to *Desulfobacteraceae* and *Desulfobulbaceae*. While population genome bins could only be resolved to the family level, recovery of 16S rRNA genes from metagenome assemblies revealed the presence of near-complete sequences related to *Desulfocapsa sulfexigens, Desulfobacula toluolea*, and an Uncultured *Desulfobacterium*. Both families (*Desulfobacteraceae* and *Desulfobulbaceae*) contain known sulfate reducers, and annotation of the genome bins revealed the metabolic potential for sulfate, sulfite, and sulfur reduction (Supplementary Text). The presence of Deltaproteobacteria in metagenome Key8 combined with the fact that produced water from Key8 was the only sample with significant amounts of sulfate (1.45 g/L) (Supplementary Table 5), suggests that these microorganisms may play a role in generating or removing sulfate.

#### Low abundance bins

Population genome bins from lower abundant taxa were recovered and include Alphaproteobacteria (*Roseovarius, Marinovum, Rhizobium*, and *Hyphomonas*), Clostridia (*Acetobacterium*), Methanomicrobia (*Methanosaeta* and *Methanospirillum*), Gammaproteobacteria (*Marinobacter*), and Epsilonproteobacteria (*Arcobacter* and *Sulfurospirillum*). In general, Alphaproteobacteria are capable of aromatic compound degradation and fermentation, and may provide the necessary substrates for methanogenesis^5,48^; The *Roseovarius* and *Hyphomonas* MAGs weren’t well resolved but contained fragments of aromatic hydrocarbon degradation pathways (Supplemental Text); *Acetobacterium* has previously been found in coalbed environments^9,10^, and is an acetogen capable of CO_2_ fixation using the Wood-Ljungdahl pathway (Supplemental Text)^49^; *Methanosaeta* spp. are acetoclastic methanogens that utilize acetate for methanogenesis; and *Marinobacter* has been found in hypersaline subsurface environments and degrade alkanes, BTEX-N, and aliphatic compounds (*e.g*., hexadecane)^50,51^.

### Intra- and inter-basin assembly-based functional profiling of Appalachian Basin metagenomes

Microbial pan-metagenomics was employed to examine the metabolic potential of the Appalachian Basin metagenomes and the functional intra-basin similarities. The functional relatedness of metagenomes (core metagenome) describes the pathways and potential driving factors common in coal environments, while the functional dissimilarity (accessory and unique metagenome) provides insight into how microbial communities are shaped by site-specific physicochemical factors. Analysis of protein sequences predicted from metagenome-assembled DNA contigs across all eight coalbed methane metagenomes, revealed a total of 429,800 predicted genes for the Appalachian Basin pan-metagenome. Genes were grouped into functional protein families based upon sequence identity (50% identity cutoff) yielding 329,564 total gene families, 52,470 accessory gene clusters, 276,720 unique gene clusters, and 374 core gene clusters (Supplementary Table 9).

The core metagenome of the Appalachian Basin represented 0.11% of the pan-metagenome, suggesting high diversity amongst microbial communities within the Appalachian coal basin. The diversity of microbial communities may be a direct result of the inherent complexity and heterogeneity of coalbed systems. Structurally coal seams are comprised of a cleat system connecting small pore spaces (<50 nm in diameter) and depending on the coal rank, have low permeability. Thus the physicochemical properties of coalbeds create micro-niches where microbial communities become physically separated and evolve into functionally distinct entities.

Annotation of the core genome revealed sequences were predominantly related to Methanomicrobiales (*Methanocalculus, Methanoregula*, and *Methanoculleus*). Other core metagenome sequences were most closely related to Methanobacteriales, Methanosarcinales, Desulfovibrionales and Pseudomonadales (Supplementary Table 10). Specifically, conserved pathways included hydrogenotrophic and acetoclastic methanogenesis, amino acid biosynthesis, and oxidative stress. Therefore, while nutrient limitation and coal heterogeneity may contribute to taxonomic and functional diversity of coal-degrading microorganisms, homogeneity of coal-degradation products (*e.g*., intermediates for methanogenesis) may play a role in the conservation of the aforementioned pathways (core metagenome).

The functional relatedness of Appalachian Basin metagenomes to other coalbed basins was also examined using pan-metagenomics. Analyses were aimed at determining how functionally related microbial communities from distinct coalbed methane wells were, and how that might infer biogenic methane production in coal from various basins. In total, fifteen metagenomes from three geographically distinct coalbed basins (three from the Alberta Basin, eight from the Appalachian Basin, and four from the San Juan Basin) were analyzed. A total of 429,182 gene clusters (pan-metagenome) were identified—23 gene families were common to all metagenomes (core metagenome), 353,745 were specific to an individual metagenome (unique metagenome), and 75,414 were found in at least one but not all metagenomes (accessory metagenome) (Supplementary Table 9).

Pan-metagenome clustering (taking into account all predicted protein sequences) of the Alberta, Appalachian, and San Juan Basin samples revealed basin-specific grouping (Figure 2), suggesting the overall metabolic potential is shaped by conditions unique to each coal region. The few exceptions were metagenome CG7 from the San Juan Basin, metagenome Trident1560D from the Alberta Basin, and metagenomes L32A and Key8 from the Appalachian Basin (Figure 2). Specifically, metagenome CG7 clustered with metagenome Key8 and both contain similar taxonomic distributions of Pseudomonadales, Clostridiales, and Bacteroidales, with little to no Archaea. The two remaining metagenomes, Trident1560D and L32A clustered together, and compared to all other coalbed methane samples contained lower abundances of Betaproteobacteria, and higher abundances of Alphaproteobacteria (Rhodobacterales and Rhizobiales). The clustering of the outliers appeared to be driven by the presence or absence of shared taxonomic groups, which is similar to trends observed by Lawson and coworkers, where habitat-specific taxonomic distribution was shaped by coal rank and physicochemical conditions exhibiting ‘patterns of endemism’ in coalbed samples^10^.

Core metagenome clustering exhibited less basin-specific groupings, but like the pan-genomic outlier clusters, aligned with relative abundances and taxonomic distributions of Bacteria and Archaea (Supplemental Figure 10). For example, formation water samples containing the highest amounts of Archaea (Methanomicrobiales) grouped together (Key7, K34, BB137, and P21), as did metagenomes comprised predominantly of Gammaproteobacteria (*Pseudomonas*) (K35, CG7, CO182 and CO183). While Basin-specific clustering in the pan-metagenome demonstrates a strong selecting force in conditions unique to each coal region, the core taxonomy-specific metagenome clustering reveals common pathways that may be taxonomically linked and evolve through exposure to physicochemical properties common to all coal environments—for example, coal-degradation products that feed into hydrogenotrophic methanogenesis pathways.

The multi-basin core metagenome contained a high proportion of gene families having KEGG or COG hits to amino acid transport metabolism, nucleotide metabolism, or xenobiotics degradation and metabolism, suggesting these pathways may be important in all coalbed environments, especially considering the exposure to hydrocarbons associated with and generated from coal. Conversely, the unique and accessory gene families were highest in cell wall/membrane/envelope biogenesis, signal transduction mechanisms, and metabolism of cofactors and vitamins. Gene families in these functional categories were presumably abundant due to the predominating physicochemical properties (impacting environmental stresses and nutrient availability) in the different basins.

Based upon pan-metagenome analysis, the unique gene families were consistently high ranging from ~78% for the Alberta basin to ~89% for the San Juan Basin, which suggests coalbed metagenomes contain a high intra-basin metabolic diversity. These findings support previous metagenomic work in hydrocarbon resource environments (oil sands, oil fields, and coalbeds), where coalbed samples were the most diverse^9^.

## DISCUSSION

Taxonomic characterization and pan-genome comparison of Appalachian Basin coalbed formation water metagenomes, combined with genome-resolved microbial metabolisms of subsurface coalbed systems, revealed functional diversity within and across coalbed basins. Specifically, high-abundance populations (Methanomicrobiales and Pseudomonadales) were highly similar (low diversity), while low-abundance populations were much more diverse--a pattern similar to other physicochemically distinct deep-sea hydrothermal vent microbial communities^52^. The high abundance of Methanomicrobiales suggests ongoing biogenic methane production in Appalachian Basin coalbeds. Furthermore, the co-occurrence of Methanomicrobiales with high salinity suggests their adaptation to saline environments. Several samples encoded for potential hydrocarbon degradation pathways, however the overall number of genes related to these pathways was limited, especially in Archaea-dominated samples. Results from this study provide a first insight of the subsurface metabolisms of Appalachian Basin coalbed microbial communities, and reveal valuable information on the potential for biogenic coal to methane conversion in the Appalachian Basin. Additional analyses (*e.g*., metatranscriptomics, metaproteomics) of coal cores with associated formation water are needed to fully assess the real-time dynamics of microbial communities in coalbed systems.

## ACKNOWLEDGEMENTS

This technical effort was performed in support of the National Energy Technology Laboratory’s ongoing research under the RES contract DE-FE0004000.

## DISCLAIMER

This report was prepared as an account of work sponsored by an agency of the United States Government. Neither the United States Government nor any agency thereof, nor any of their employees, makes any warranty, express or implied, or assumes any legal liability or responsibility for the accuracy, completeness, or usefulness of any information, apparatus, product, or process disclosed, or represents that its use would not infringe privately owned rights. Reference therein to any specific commercial product, process, or service by trade name, trademark, manufacturer, or otherwise does not necessarily constitute or imply its endorsement, recommendation, or favoring by the United States Government or any agency thereof. The views and opinions of authors expressed therein do not necessarily state or reflect those of the United States Government or any agency thereof.

